# Sobp modulates Six1 transcriptional activation and is required during craniofacial development

**DOI:** 10.1101/2021.04.05.438472

**Authors:** Andre L. P. Tavares, Karyn Jourdeuil, Karen M. Neilson, Himani D. Majumdar, Sally A. Moody

## Abstract

Branchio-oto-renal syndrome (BOR) is a disorder characterized by hearing loss, craniofacial and/or renal defects. Mutations in the transcription factor Six1 and its cofactor Eya1, both required for otic development, are linked to BOR. We previously identified Sobp as a potential Six1 cofactor and *SOBP* mutations in mouse and humans cause otic phenotypes; therefore, we asked whether Sobp interacts with Six1 and thereby may contribute to BOR. Co-IP and immunofluorescence experiments demonstrate that Sobp binds to and co-localizes with Six1 in the cell nucleus. Luciferase assays show that Sobp represses Six1+Eya1 transcriptional activation. Experiments *in Xenopus* embryos that either knockdown or increase expression show that Sobp is required for formation of ectodermal domains at neural plate stages. In addition, altering Sobp levels disrupts otic vesicle development and causes craniofacial cartilage defects. Expression of *Xenopus* Sobp containing the human mutation disrupts the pre-placodal ectoderm similar to full-length Sobp, but other changes are distinct. These results indicate that Sobp modifies Six1 function, is required for vertebrate craniofacial development, and identifies Sobp as a potential candidate gene for BOR and other deafness syndromes.

**Summary statement:** Sobp interacts with Six1 in the cell nucleus and represses the Six1+Eya1 transcriptional activation. In *Xenopus* embryos, Sobp functions during early stages of inner ear development.

## INTRODUCTION

Hearing is an important component of human communication and hearing loss disorders negatively affect quality of life. Around 32 million children worldwide have disabling hearing loss, 40% of which have a genetic cause (Krug, 2016). Although hundreds of genes are associated with syndromic and non-syndromic congenital hearing loss, many cases still have an unknown cause (Shearer et al., 2017). Branchio-otic and branchio-oto-renal syndromes (BOR) are autosomal dominant disorders in which affected individuals present with variable degrees of hearing loss due to defects in inner, middle and outer ears, as well as branchial arch-associated dysmorphologies including branchial fistulas and cysts (Moody et al., 2015, Smith, 2018). Mutations in *EYA1* or *SIX1* have been identified in approximately 50% of affected individuals, but the underlying genetic causes of remaining cases are unknown (Ruf et al., 2003, Ruf et al., 2004, Sanggaard et al., 2007, Lee et al., 2007, Klingbeil et al., 2017). Thus, identifying genes that interact with SIX1 may uncover novel genes linked to BOR or other deafness syndromes.

The Six1 transcription factor, homologous to *Drosophila* Sine oculis (So) (Cheyette et al., 1994), plays a role in many cellular processes and in the embryo regulates the development of craniofacial tissues affected in BOR (Kawakami et al., 2000, Kumar, 2009, Xu, 2013, Moody and LaMantia, 2015). Loss of Six1 in mouse and *Xenopus* causes several craniofacial defects including disruption of inner, middle and outer ear development (Zheng et al., 2003, Li et al., 2003, Laclef et al., 2003, Ozaki et al., 2004, Brugmann et al., 2004, Schlosser et al., 2008, Guo et al., 2011, Tavares et al., 2017, Sullivan et al., 2019). A hallmark of Six1 transcriptional activity is that it can be modulated by co-factors that influence it to either activate or repress target genes (Ohto et al., 1999, Li et al., 2003, Silver et al., 2003, Brugmann et al., 2004). Transcriptional activation is mediated by the Eya family of co-activators. Upon binding to Six1, Eya factors are translocated into the nucleus where Eya phosphatase activity triggers the recruitment of co-activators that switch Six1 function from repression to activation (Ohto et al., 1999, Li et al., 2003).

A screen to identify proteins that interact with *Drosophila* So identified So binding protein (Sobp), which is co-expressed with *so* in the anterior region of the eye disc (Kenyon et al., 2005a). In mouse, *Sobp* was identified as the spontaneous recessive mutation causing deafness and vestibular-mediated circling behavior in the Jackson circler (*jc*) mouse (Calderon et al., 2006, Chen et al., 2008). In humans, *SOBP* was linked to mental retardation, anterior maxillary protrusion, strabismus and mild hearing loss (MRAMS; OMIM #613671) in 7 individuals of the same family. The homozygous mutation inserts an early stop codon at arginine 661 (R661X) causing a 212 amino acid truncation in the C-terminus (Basel-Vanagaite et al., 2007, Birk et al., 2010).

Since *Sobp* mutations cause otic phenotypes in mouse and humans, and is expressed in the pre-placodal ectoderm (PPE) and otic vesicle in *Xenopus* embryos (Neilson et al., 2010), we speculated that it may function by associating with Six1 and thereby contribute to BOR (Moody et al., 2015). Herein we address the developmental function of Sobp. We found that *sobp* is expressed in the same tissues as *six1*, but in complementary domains suggesting it may repress Six1 activity. We show that Sobp binds to Six1, competes with Eya1 binding to the complex and significantly reduces Six1+Eya1 transcriptional activation. Structural and functional analyses show that Sobp constructs either lacking a conserved C-terminal nuclear localization signal (NLS) or containing a mutation similar to human R661X (*Xenopus* Sobp R651X) can still interact with Six1, indicating that the domain that interacts with Six1 is not in the C-terminus. *In vivo* studies establish Sobp as a critical factor for PPE and neural crest (NC) gene expression and later otic vesicle development. These findings also show that while the R651X mutation disrupts gene expression in the PPE in the same manner as full-length Sobp, changes in other domains and otic vesicle patterning are distinct. Together, these results demonstrate that Sobp plays a critical role in Six1 transcriptional function during several aspects of vertebrate craniofacial development, and suggest it may be a candidate gene for BOR and other deafness syndromes.

## RESULTS

### Sobp expression in comparison to that of Six1

We reported previously that *sobp* is expressed in several of the embryonic tissues that also express *six1*, including the PPE and otic vesicle (Neilson et al., 2010). A closer comparison at the neural plate stage showed that *sobp* expression was stronger in the anterior PPE and weaker in the posterior PPE, whereas *six1* expression was the reverse (Fig. 1A-B). At larval stages both *six1* and *sobp* were expressed in the medial wall of the otic vesicle, but *sobp* expression appeared more intense dorsally compared to its ventral domain (Fig. 1C-D; Fig. S1A-B). Co-transfection assays in HEK293T cells showed that Sobp protein was mainly nuclear (Fig. 1E-G), as previously described in mouse (Chen et al., 2008), where it co-localized with Six1 in the majority of cells expressing both constructs (Fig. 1H-K). These findings are the first direct evidence in vertebrates of Sobp and Six1 co-expression in tissues and cellular compartments that would allow their interaction.

**Figure 1.**
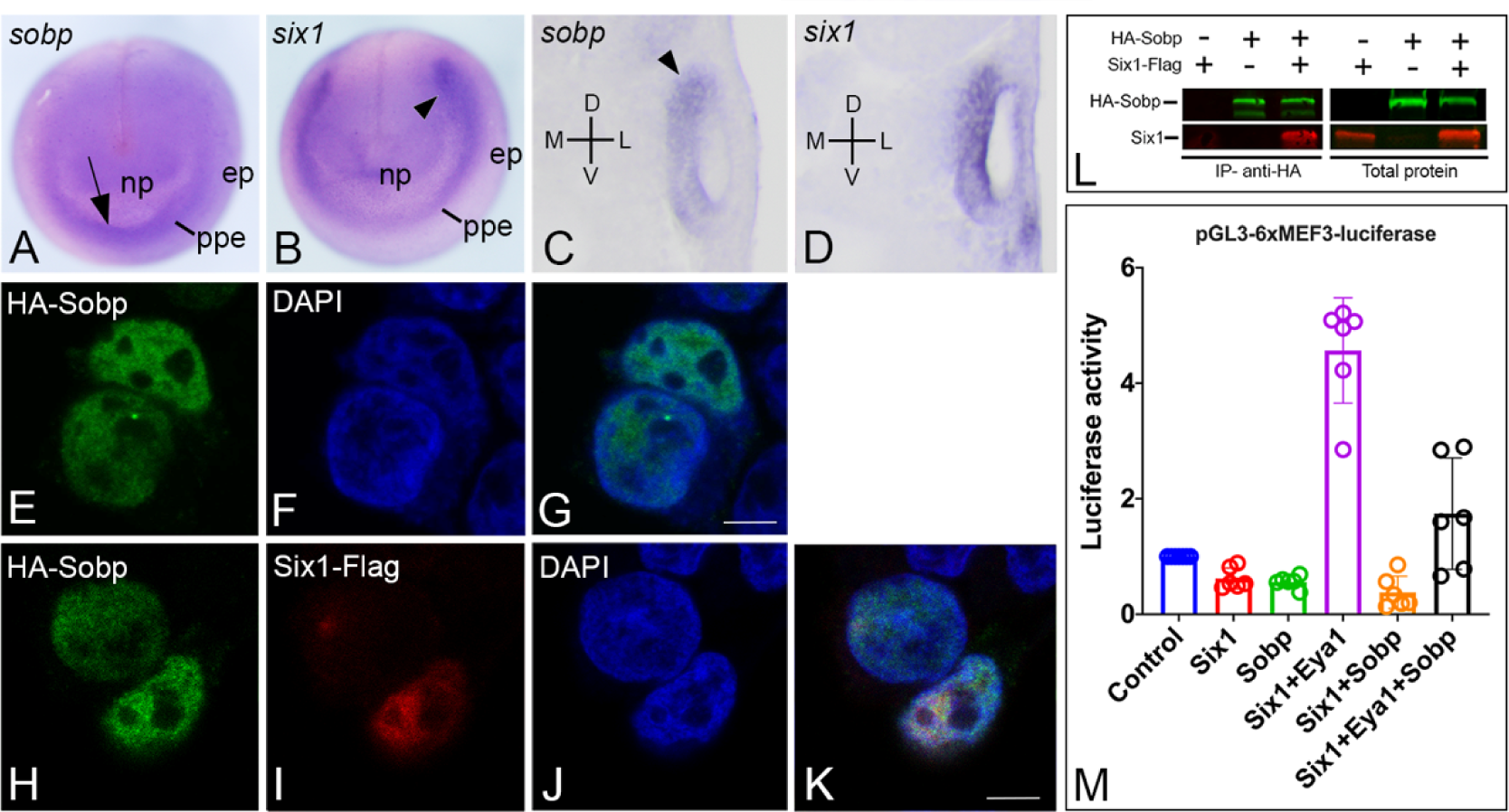
Sobp is expressed with Six1 in the cell nucleus and represses its transcriptional activation. **A-D.** In situ hybridization for *sobp* (A, C) and *six1* (B, D). At neural plate stages (**A-B**), while *sobp* expression in the PPE overlaps with that of *six1*, its expression is more intense in the anterior domain (arrow) whereas *six1* expression is more intense in the posterior domain (arrowhead). In transverse sections through the larval otic vesicle (**C-D**), *sobp* is expressed with *six1* in the ventral-medial wall. Note that *sobp* expression is more intense at the dorsal pole (arrowhead). D, dorsal; ep, epidermis; L, lateral; M, medial; np, neural plate; ppe, pre-placodal ectoderm; V, ventral. **E-K**. Confocal images of HEK293T cells expressing HA-Sobp (green, E-G) and cells co-expressing HA-Sobp (green) and Six1-Flag (red) (H-K). Sobp is localized in the cell nuclei in both the absence and presence of Six1-Flag. Cell nuclei are stained with DAPI (blue, F, G, J, K). Bars: 5μm. **L.** HEK293T cells were co-transfected with combinations of HA-Sobp and/or Six1-Flag followed by multiplex fluorescence Western blot detection for HA-Sobp (green) and Six1-Flag (red). Six1 was detected after HA-Sobp was immunoprecipitated with anti-HA magnetic beads (IP, left two rows). Right two rows show expression of the constructs prior to immunoprecipitation. **M**. Graph depicting the luciferase activity of the pGL3-6xMEF3-luciferase reporter in HEK293T cells transfected with different combinations of constructs expressing Six1, Eya1 and/or Sobp. Data are normalized to Renilla expressed with a constitutive promoter. Luciferase activity is significantly induced (p<0.0001) by Six1+ Eya1, whereas Sobp reduces this induction to levels indistinguishable from control (Six1+Eya1 vs. Six1+Eya1+Sobp: p<0.0001; Control vs. Six1+Eya1+Sobp: p=0.2226). Neither Six1 (p=0.9984), Sobp (p=0.9184) nor Six1+Sobp (p=0.9184) caused significant changes in luciferase activity compared to control. Experiments were repeated in duplicate at least 3 independent times. Error bars represent standard deviation (SD) with circles depicting individual data points.

### Sobp binds to Six1 and modulates Six1+Eya1 transcriptional activation

*Drosophila* Sobp was shown to bind to So by yeast two hybrid and GST-pulldown assays (Kenyon et al., 2005a). To test if this interaction occurred with vertebrate proteins, HEK293T cells were co-transfected with constructs driving expression of Six1-Flag and/or HA-Sobp followed by immunoprecipitation (IP). Western-blot analysis detected Six1 after HA-Sobp IP (Fig. 1L); reverse IP confirmed this finding (data not shown).

To assess whether Sobp modulates Six1 transcriptional activity, HEK293T cells were co-transfected with a Six1-inducible reporter (Ford et al., 2000) and different combinations of Six1, Eya1 and/or Sobp (Fig. 1M; Fig. S2). As demonstrated previously (Patrick et al., 2009), in the presence of Eya1, Six1 induced a significant increase in luciferase activity over control (∼5 fold increase, p<0.0001). In contrast, Sobp by itself (p=0.8899) or in the presence of Six1 (p=0.9184) did not cause a significant change in luciferase activity. However, co-transfection of Sobp with both Six1 and Eya1 significantly repressed luciferase activity compared to Six1+Eya1 levels (p<0.0001), returning luciferase activity to levels statistically indistinguishable from control (p=0.2226). These data show that Sobp can be classified as a bona fide Six1 co-factor because it binds to Six1 and is able to significantly interfere with Six1+Eya1 transcriptional activation.

Pa2g4 repressed Six1+Eya1 transcriptional activity by competing with the ability of Eya1 to bind to Six1 (Neilson et al., 2017). To verify whether Sobp also competes for binding to Six1, HEK293T cells were co-transfected with Six1-Flag, Myc-Eya1 and increasing amounts of HA-Sobp. Co-IP experiments showed that equimolar amounts (1.0X) of Six1, Eya1 and Sobp did not disrupt Six1 binding to Eya1, whereas the amount of Six1 bound to Eya1 was diminished at a 5-fold or 10-fold increase in Sobp (Fig. 2A). Because Eya factors are localized in the cytosol and become nuclear when bound to Six factors (Ohto et al., 1999, Li et al., 2003) (Fig. S3A-B, D-E, G-H, J-K), we tested whether the competition for binding to Six1 would disturb the ability of Six1 to translocate Eya1 to the nucleus. Cells co-transfected with combinations of Six1-Flag, Myc-Eya1 and increasing concentration of HA-Sobp showed that at equimolar amounts of Six1, Eya1 and Sobp, Eya1 was translocated to the nucleus by Six1 (Fig. 2B, F, J, N) with most transfected cells also expressing Sobp in the nucleus (Fig. 2C, G, K, O).

**Figure 2.**
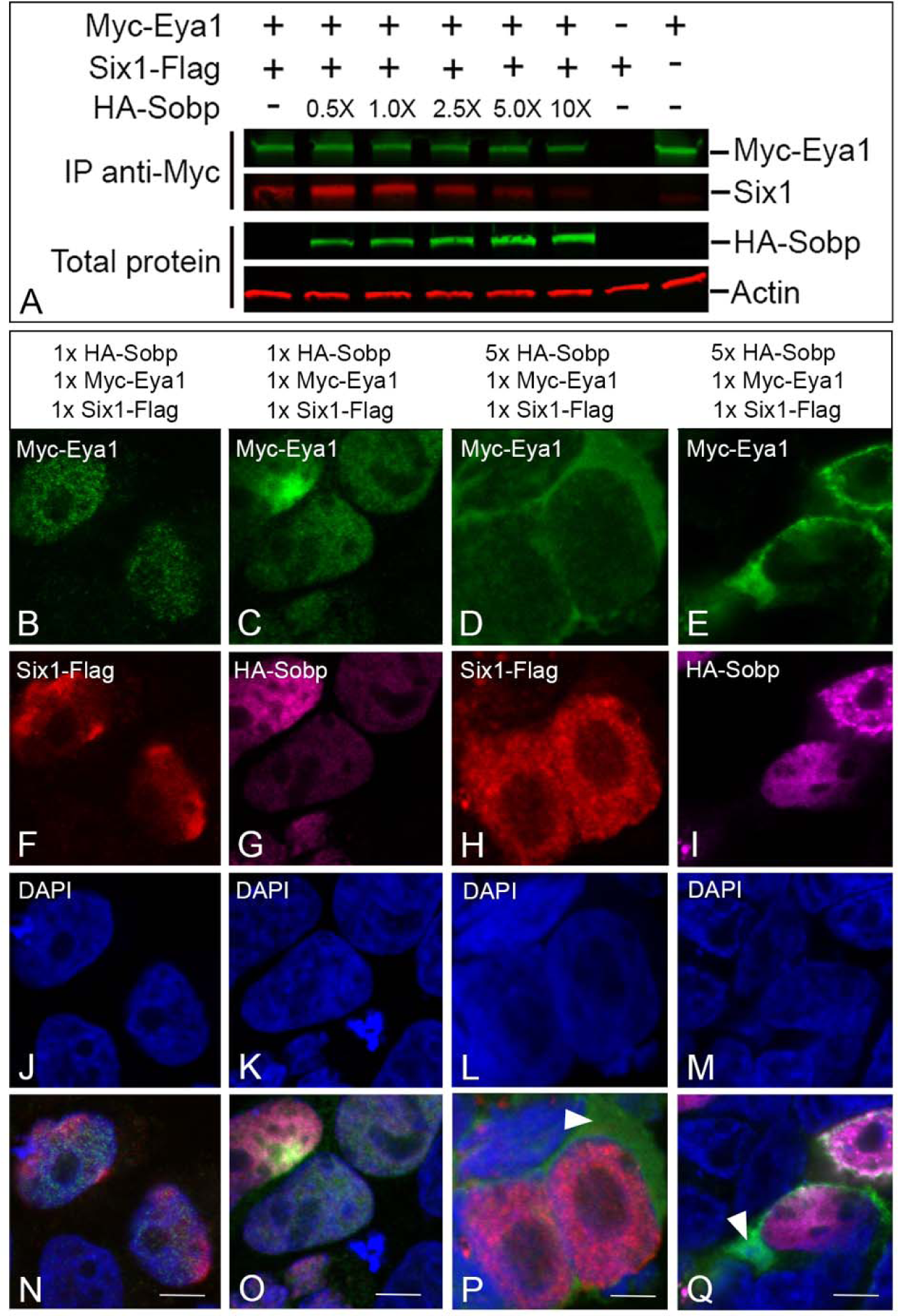
Sobp reduces Six1+Eya1 transcriptional activation by disrupting the Six1/Eya1 interaction. **A.** HEK293T cells co-transfected with equimolar amounts of Six1-Flag and/or Myc-Eya1 were additionally transfected with increasing amounts of HA-Sobp. The amount of Six1 bound to Myc-Eya1decreased with increasing levels of HA-Sobp (2.5x, 5.0x, 10x). The two bottom rows show expression before immunoprecipitation of increasing levels of HA-Sobp and β-actin as loading control . **B-Q.** Confocal images of HEK293T cells expressing Myc-Eya1 (green, B-E, N-Q), Six1-Flag (red, F, H, N, P) and HA-Sobp (magenta, G, I, O, Q). Myc-Eya1 was completely translocated into the cell nucleus by Six1 when cells received equimolar amounts (1X) of Six1-Flag, Myc-Eya1 and HA-Sobp (B-C, F-G, N-O), whereas cytosolic Myc-Eya1 (arrow in P and Q) was detected in many cells when there was a 5-fold increase in HA-Sobp. Nuclear DAPI staining, blue (J-Q). Bars: 5 μm.

Remarkably, Sobp was able to bind to (Fig. S3M) and partially translocate Eya1 to the nucleus (Fig. S3C, F, I, L) without Six1 co-expression. However, and in agreement with the competition Co-IP results, a 5-fold increase in Sobp led to detection of Eya1 in the cytosol (Fig. 2D-E, P-Q) even though both Six1 and Sobp were nuclear (Fig. 2H-I, L-M). Together, these data indicate that Sobp repression of Six1+Eya1 transcriptional activation is achieved through a dose-dependent competition mechanism.

### Six1 can transport Sobp to the nucleus

Because Sobp is located in the nucleus in the absence of Six1 co-transfection (Fig. 1E-G) (Chen et al., 2008), it must contain a nuclear localization signal (NLS). A bioinformatic analysis (Kosugi et al., 2009) identified in *Xenopus* Sobp N-terminal and C-terminal putative nuclear localization signals (NLS), similar to what has been detected in the mouse (Chen et al., 2008) and human (Birk et al., 2010) sequences. However, only the putative C-terminal NLS had a cutoff score above 8, which indicates protein localization exclusively in the nucleus (Fig. 3A-B) (Kosugi et al., 2009). Deletion of this C-terminal domain caused HA-Sobp-NLSdel to be detected mainly in the cytosol (Fig. 3C-E). Interestingly, when cells additionally expressed Flag-tagged Six1, Sobp-NLSdel was translocated to the nucleus (Fig. 3F-I). Removal of the NLS did not disrupt binding of Sobp to Six1 (Fig. 3J) or diminished the ability of Sobp to reduce Six1+Eya1 transcriptional activation (p<0.0001; Fig. 3K). These findings identify a single C-terminal NLS in Sobp and demonstrate that in its absence Sobp can bind to Six1 in the cytosol and be translocated to the nucleus via this interaction.

**Figure 3.**
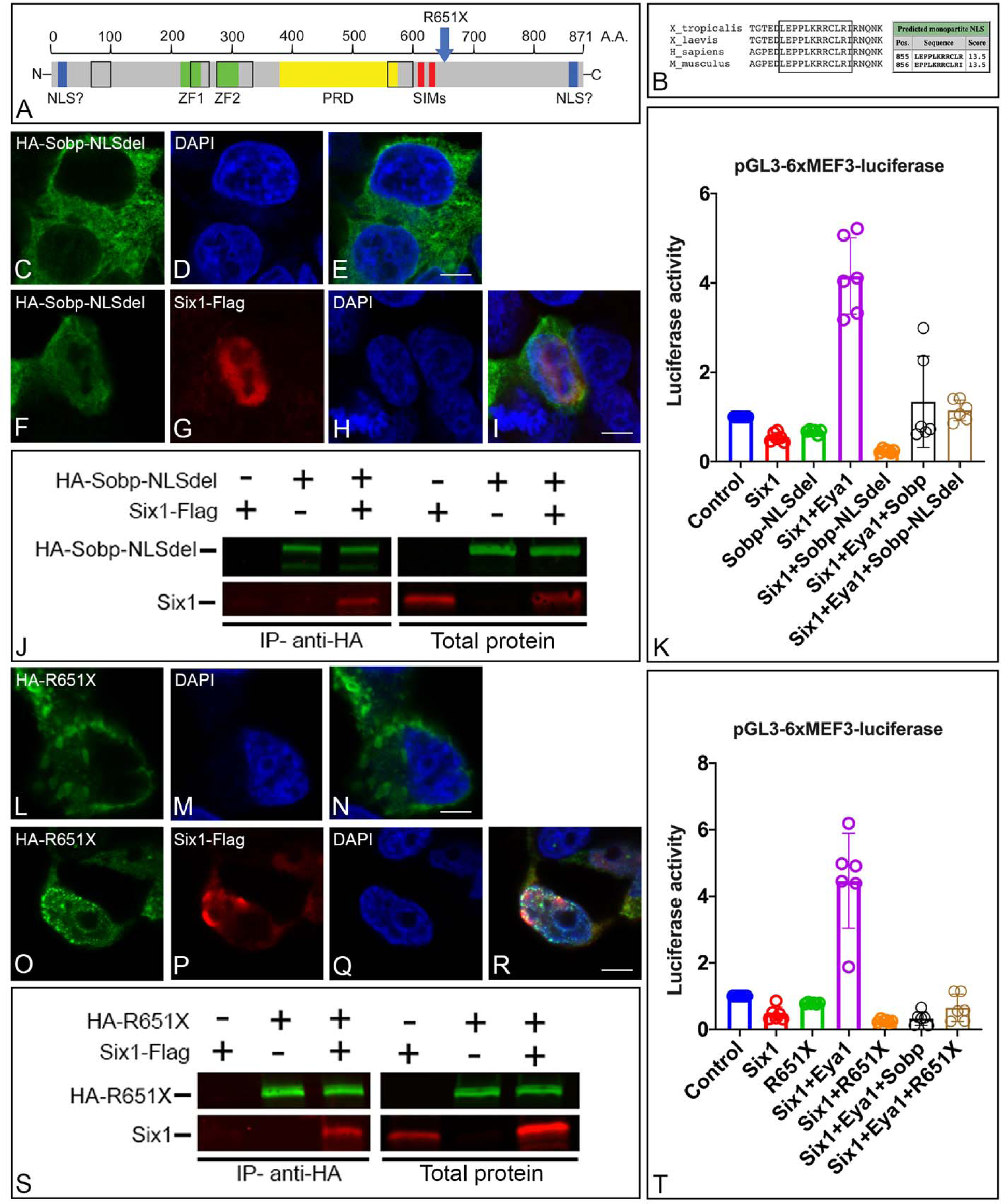
Deletion of the nuclear localization signal and the R651X mutation do not disrupt Sobp interaction with Six1. **A**. Schematic representation of the Sopb protein structure showing its different domains and the location of the R651X mutation identified in MRAMS patients. Black empty boxes denote protein domains that are highly conserved between different species including *D. melanogaster*, *M. musculus*, *G. gallus* and *X. laevis/tropicalis*; blue boxes/NLS denote putative nuclear localization signals; green boxes/ZF1/ZF2 denote FCS zinc finger domains; yellow box/PRD denotes proline rich domain; red boxes/SIMs denote SUMO interacting motifs. **B.** Comparison of the C-terminal region of Sobp between species showing a highly conserved domain that is predicted to be a NLS with a high cutoff score according to cNLS mapper. **C-I.** Confocal images of HEK293T cells expressing a construct lacking the C-terminal NLS (HA-Sobp-NLSdel, green). This construct is cytosolic in the majority of the transfected cells (C, E). Cells co-expressing HA-Sobp-NLSdel with Six1-Flag (red, G, I) show partial translocation to the nucleus (green, F, I). DAPI (blue, D, E, H, I). Bars: 5 μm. **J.** HEK293T cells were co-transfected with combinations of HA-Sobp-NLSdel and/or Six1-Flag followed by multiplex fluorescent Western blot detection for HA-Sobp-NLSdel (green) and Six1 (red). Six1 was detected after HA-Sobp-NLSdel was immunoprecipitated with anti-HA magnetic beads (IP, left two rows). Right two rows show expression of the constructs before immunoprecipitation. **K**. Graph depicting the luciferase activity of the pGL3-6xMEF3-luciferase reporter in HEK293T cells transfected with different combinations of constructs expressing Six1, Eya1, Sobp and/or Sobp-NLSdel. The C-terminal NLS is not required for repression of Six1+Eya1 transcriptional activation (Control vs Six1+Eya1+Sobp-NLSdel, p=0.9959; Six1+Eya1+Sobp vs Six1+Eya1+Sobp-NLSdel, p=0.9912). Experiments were repeated in duplicate at least 3 independent times. Error bars represent SD with circles depicting individual data points. **L-R.** Confocal images of HEK293T cells expressing the R651X mutant (HA-R651X, green). The mutant is cytosolic in the majority of transfected cells (L, N). Cells co-expressing R651X and Six1-Flag (red, P, R) show partial translocation of HA-R651X to the cell nucleus (green, O, R). Cell nuclei are stained with DAPI (M, N, Q, R). Bars: 5 μm. **S.** HEK293T cells were co-transfected with combinations of HA-R651X and/or Six1-Flag followed by multiplex fluorescence Western blot detection for HA-R651X (green) and Six1 (red). Six1 was detected after HA-R651X was immunoprecipitated with anti-HA magnetic beads (IP, left two rows). Right two rows show expression of the constructs before immunoprecipitation. **T.** Graph depicting the luciferase activity of the pGL3-6xMEF3-luciferase reporter in HEK293T cells transfected with different combinations of constructs expressing Six1, Eya1, Sobp and/or R651X. Truncated Sobp (R651X) is still able to repress Six1+Eya1 transcriptional activation (Control vs Six1+Eya1+R651X, p=0.8603; Six1+Eya1+Sobp vs Six1+Eya1+R651X, p=0.9362). Replication of experiments and error bars as in Figure 1.

### The human R661X mutation does not disrupt its interaction with Six1

We introduced the human R661X mutation found in a MRAMS family (Birk et al., 2010) in *Xenopus* Sobp by replacing the arginine located at amino acid 651 with a stop codon (c.1951 A>T; p.R651X) generating a truncated Sobp lacking part of the C-terminus including the NLS (Fig. 3A). As expected, HA-R651X located primarily in the cytosol (Fig. 3L-N) since the NLS was missing. Cytosolic HA-R651X was translocated to the nucleus in most cells also expressing Six1-Flag (Fig. 3O-R). Despite the large deletion, R651X still bound to Six1 (Fig. 3S) and significantly reduced Six1+Eya1 transcriptional activation (p<0.0001; Fig. 3T). These data demonstrate that although the R651X mutation lacks a large portion of the C-terminus the domain for Six1 interaction remains intact.

### Sobp is required for development of neural crest and placode domains

To determine if Sobp has a role in NC and PPE development, we performed two types of knock-down (KD) experiments: a translation-blocking antisense morpholino oligonucleotide (MO) against *sobp* and an F0 analysis after CRISPR/Cas9-mediated genome editing (Figure 4A-H, Q-R, U; Fig. S4). These embryos were assessed for the expression of genes that mark the neural plate (*sox2*), NC (*foxd3*), PPE (*six1*) or epidermis (*krt12.4*) at neural plate stages. Sobp KD by MO or by CRISPR each caused an expansion of the *sox2* domain (MO: 77.0%, Fig. 4A, Q; CRISPR: 64.9%, Fig. 4E, R) and reduction of the *foxd3* (MO: 90.2%, Fig. 4B, Q; CRISPR: 66.6%, Fig. 4F, R), *six1* (MO: 73.9%, Fig. 4C, Q; CRISPR: 76%, Fig. 4G, R) and *krt12.4* (MO: 95.8%, Fig. 4D, Q; CRISPR: 78.6%, Fig. 4H, R) domains on the injected side of the embryos. qPCR analysis at the same stage (Fig. 4U) showed decreases in mRNA levels for *foxd3* (MO: p<0.001/q=0.000602; CRISPR: p<0.01/q=0.002520) and *six1* (MO: p<0.0001/q=0.000007; CRISPR: p<0.001/q=0.000274), but not for *sox2* (MO: p=0.033437/q=0.033771; CRISPR: p=0.071790/q=0.036254) or *krt12.4* (MO: p=0.114167/q=0.086482; CRISPR: p=0.104590/q=0.042254); the discrepancy of the latter two genes from ISH analyses is likely due to transcripts from whole embryos being analyzed. We also monitored levels of *sobp* mRNA (Fig. 4U) and confirmed that, as expected, it was decreased by CRISPR/Cas9 editing likely followed by nonsense-mediated mRNA decay (p<0.0001/q=0.000162), but not by a translation-blocking MO (p=0.359946/q=0.218127). To ensure MO specificity, embryos were injected with Sobp MO plus an MO-insensitive *sobp* mRNA and assessed for *foxd3* expression (Fig. 4B, Q). Only 29.4% of these embryos (*cf*. 90.2% MO alone) had reduced *foxd3* expression (Fig.S4C-D), demonstrating significant rescue.

**Figure 4.**
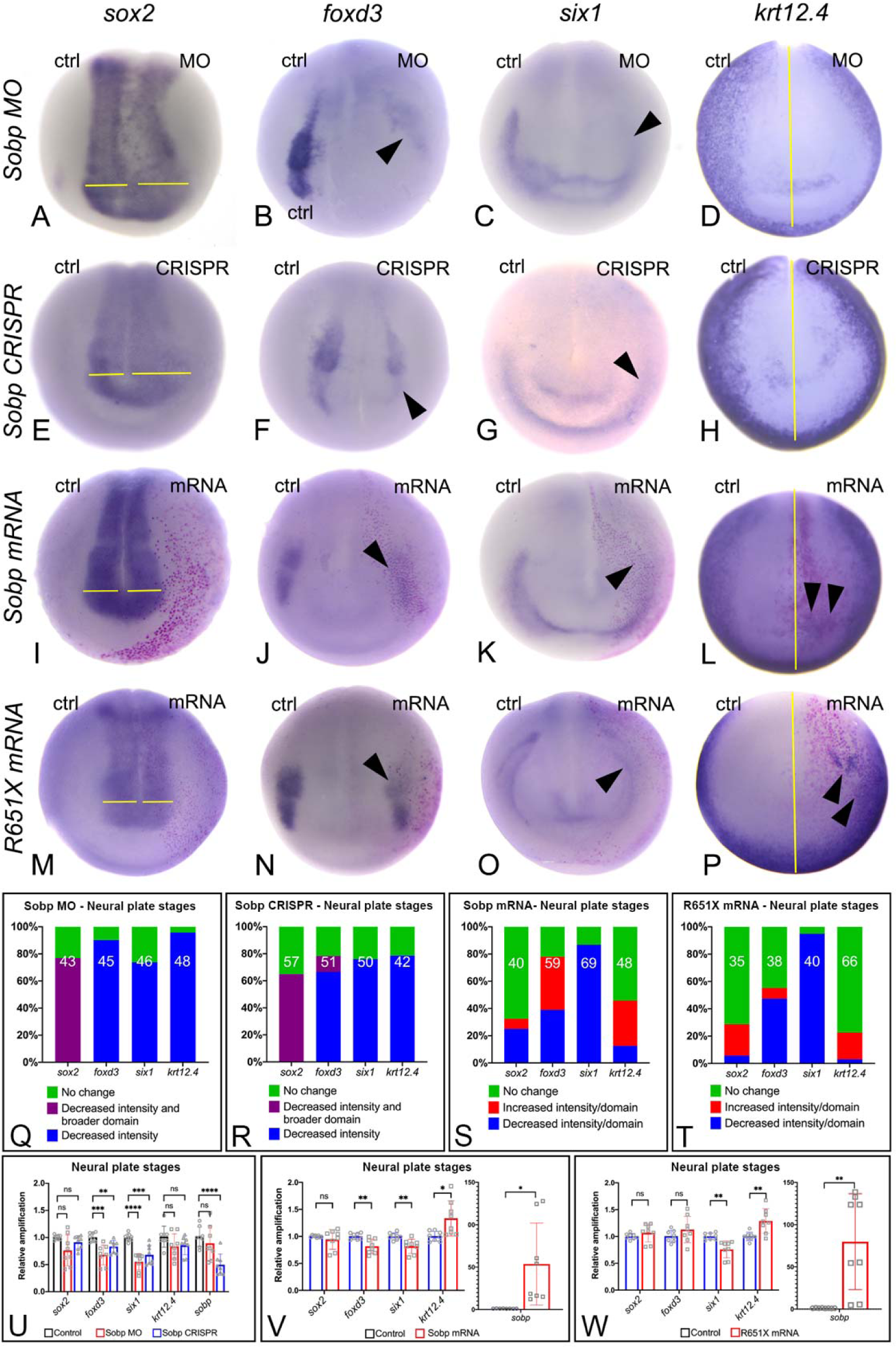
Sobp is required for proper formation of the embryonic ectodermal domains at neural plate stages. **A-P.** In situ hybridization for *sox2* (neural plate), *foxd3* (neural crest), *six1* (PPE) and *krt12.4* (epidermis). Images are representative of the most frequent phenotype except for *krt12.4* in L and P. (**A-H**) Knock-down of Sobp on one side in morphants (MO) or F0 crispants (CRISPR) reduced the intensity of *sox2* expression concomitant with expansion of its domain, indicated by yellow lines (A, E). They also reduced expression of *foxd3* (arrowhead, B, F), *six1* (arrowhead C, G) and *krt12.4* (indicated by distance from the midline, yellow line, D,H). Increased expression of Sobp (I-L) or the R651X mutant (M-P) on one side caused a decrease in the expression of *foxd3* (arrowhead, J, N) and *six1* (arrowhead, K, O), whereas *sox2* expression was unchanged (I, M). Although *krt12.4* expression was unchanged in most embryos, we detected ectopic expression overlapping the lineage tracer (arrowheads, L, P). **Q-T.** Frequencies of changes in gene expression illustrated in panels A-P. The number in each bar denotes sample sizes. **U.** qPCR analysis of whole neural plate embryos injected with MO or after CRISPR shows that Sobp KD caused a significant decrease in the mRNA levels for *foxd3* (MO: ∼33%; CRISPR: ∼20%) and *six1* (MO: ∼40%; CRISPR: ∼32%) relative to uninjected control embryos, whereas changes in *sox2* or *krt12.4* were not significant. Levels of *sobp* mRNA verified reduced transcripts after CRISPR (MO: not significant; CRISPR: ∼50% decrease). **V.** qPCR analysis of whole embryos injected with *sobp* mRNA shows that increasing Sobp (∼50-fold increase) significantly reduced mRNA levels for *foxd3* (∼20%) and *six1* (∼20%), and increased that of *krt12.4* (∼1.3 fold), whereas *sox2* levels were not significantly affected. **W.** qPCR analysis of embryos injected with *R651X* mRNA shows that increased expression (∼80-fold increase) significantly reduced mRNA levels for *six1* (∼25%) and increased that of *krt12.4* (∼1.3 fold), whereas *foxd3* and *sox2* levels were not significantly affected. ns. not significant, * p<0.05, ** p<0.01, *** p<0.001, **** p<0.0001. qPCR experiments were repeated at least 4 independent times. Error bars represent SD with symbols depicting individual data points.

To determine whether increasing Sobp protein above endogenous levels altered gene expression, we targeted *sobp* mRNA to the dorsal-animal and ventral-animal blastomeres of 8-cell embryos (Fig. 2I-L, S, V), which are major precursors of NC and PPE (Moody and Kline 1990). Even though we detected a decrease in the *sox2* domain in some embryos (25.0%), the majority showed no changes on the injected side (67.5%, Fig. 4I, S). The effects on *foxd3* were pleiotropic: the domain was smaller in 39.0% (Fig. 4J, S) and broader in 39.0% (Fig. 4S) of embryos. The *six1* domain was smaller in the majority of the embryos (86.9%, Fig. 4K, S). Even though the *krt12.4* domain was increased in some embryos with ectopic expression of this gene overlapping the lineage tracer (33.3%, Fig. 4L, S), this marker was unchanged in most embryos (54.2%, Fig. 4S). qPCR analysis at the same stage (Fig. 4V) showed that increasing Sobp significantly reduced mRNA levels of *foxd3* (p<0.01/q=0.002780) and *six1* (p<0.01/q=0.002780), and increased levels of *krt12.4* (p<0.05/q=0.009927); *sox2* levels were not significantly affected (p=0.397516/q=0.200746). These data demonstrate that altering the levels of Sobp disrupts the relative sizes and gene expression pattern of the embryonic ectodermal domains.

Because the R651X mutation does not disrupt the interaction of this protein with Six1, we assessed whether R651X would cause the same changes in genes expression as full-length Sobp. The *sox2* domain was not affected in the majority (71.4%, Fig. 4M, T) of embryos. The *foxd3* domain was reduced in about half (47.4%, Fig. 4N, T) and increased in a small number of embryos (7.9%, Fig. 4T). The *six1* domain was decreased at high frequency (95.0%, Fig. 4O, T) and the *krt12.4* domain was unchanged in most embryos (77.3%, Fig. 4T); a low percentage showed a slight increase with ectopic expression of *krt12.4* in the area where the lineage tracer was present (19.7%, Fig. 4P, T). A comparison of mutant versus full-length Sobp phenotype frequencies showed that *sox2* expression was increased significantly more frequently (p<0.05), *foxd3* (p<0.01) and *krt12.4* (p<0.05) were increased less frequently, in embryos expressing R651X, whereas effects on *six1* were indistinguishable (p=0.3772). qPCR analysis of whole embryos (Fig. 4W) showed a significant decrease in *six1* (p<0.01/q=0.002134) and significant increase in *krt12.4* (p<0.01/q=0.004706), but no significant changes in *foxd3* (p=0.211730/q=0.142565) or *sox2* (p=0.307810/q=0.155444) mRNA levels. These data demonstrate that introduction of the R651X mutation in Sobp did not alter changes in *six1* expression caused by full-length Sobp, but showed subtle changes in effects on *sox2*, *foxd3* and *krt12.4* expression. Importantly, these results indicate that except for its ability to translocate to the nucleus, R651X is sufficiently functional to alter early ectodermal gene expression.

### Proper levels of Sobp are required for otic vesicle development

Since both KD and increased expression of Sobp altered PPE gene expression, we analyzed whether these changes led to disruptions in the otic vesicle at larval stages (Fig. 5). KD of *sobp* by CRISPR (Fig. 5A, D, G, J, M) revealed a decrease in the intensity and domain size of the otic expression of *dlx5* (42.9%, Fig. 5D, J) and *pax2* (54.0%, Fig. 5G, J); interestingly, *six1* was not altered in most *sobp* crispants (68.6%, Fig. 5A, J) as only 25.7% of embryos showed a slight decrease in expression (Fig. 5J). Similar findings were shown by qPCR (Fig. 5M): decreased *dlx5* (p<0.01/q=0.002700) and *pax2* (p<0.01/q=0.002668) and no significant change in *six1* (p=0.450307/q=0.227405).

**Figure 5.**
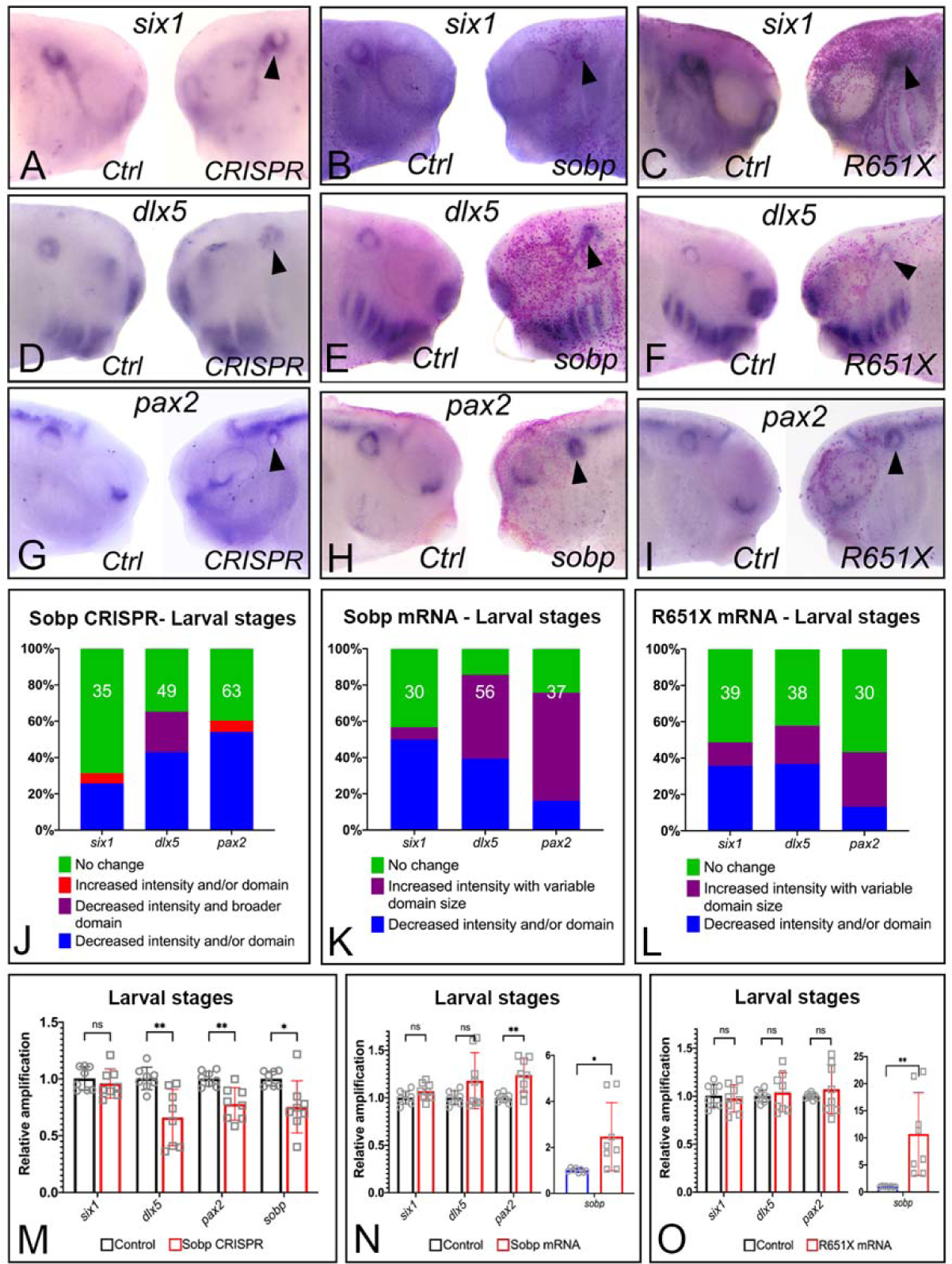
Dorsal-ventral patterning of the otic vesicle requires proper expression levels of Sobp. **A-I.** In situ hybridization for *six1* (A-C), *dlx5* (D-F) and *pax2* (G-I) at larval stages. Images in each box are the control and injected sides of the same embryo, and are representative of the most frequent phenotype for CRISPR (A, D, G) and *sobp* mRNA (C, F, I). Sobp KD leads to decreased otic expression of *dlx5* (arrowhead, D) and *pax2* (arrowhead in G), whereas *six1* expression is unchanged (arrowhead, A). Increased Sobp causes a decrease in *six1* expression (arrowhead, B) and increased expression with a variable domain size of *dlx5* (arrowhead, E) and *pax2* (arrowhead, H). Less frequently, increased R651X expression caused a decrease in *six1* (arrowhead, C) and *dlx5* (arrowhead, F) and increased expression of *pax2* (arrowhead, I). **J-L.** Frequencies of changes in gene expression illustrated in panels A-I. The number in each bar denotes the sample size. **M.** qPCR analysis of larval heads after CRISPR (∼25% decrease in *sobp* mRNA) shows significant decrease in *dlx5* (∼34%) and *pax2* (∼22%) mRNAs, whereas changes in *six1* are not significant. **N.** qPCR analysis of larval heads injected with *sobp* mRNA (∼2.5-fold increase) shows a significant increase in *pax2* (∼1.4 fold); changes in *six1* and *dlx5* are not significant. **O.** qPCR analysis of larval heads injected with *R651X* mRNA (∼10-fold increase) shows no significant changes in *six1*, *dlx5* or *pax2*. ns. not significant, * p<0.05, ** p<0.01, *** p<0.001, **** p<0.0001. qPCR experiments were repeated at least 4 independent times. Error bars represent SD with symbols depicting individual data points.

Increased Sobp had variable effects on otic vesicle genes (Fig. 5B, E, H, K, N). *six1* expression was decreased in 50.0%, increased in 6.7% and unchanged in 43.3% of larvae (Fig. 5B, K). *dlx5* showed increased expression with a variable domain size in 46.4% and decreased expression and domain size in 39.3% of larvae (Fig. 5E, K). *pax2* showed increased expression with a variable domain size in 59.5% and decreased expression in 16.2% of larvae (Fig. 5H, K). qPCR of whole heads confirmed the effects were variable because there were no significant changes in the mRNA levels for *six1* (p=0.106946/q=0.079244) or *dlx5* (p=0.117690/q=0.079244) (Fig. 5N). Only *pax2* showed a significant increase in mRNA levels (p<0.01/q=0.005414) after increased Sobp expression. These findings show that altering Sobp levels that change PPE gene expression are followed by disruptions of otic vesicle development.

We tested whether the R651X mutation of *sobp* also altered otic gene expression. Effects on otic gene expression were variable with most of the analyzed larvae showing no changes in *six1* (51.3%), *dlx5* (42.1%) or *pax2* (56.7%; Fig. 5L). Less frequently, we observed a decrease in the expression and domain size for *six1* (35.9%, Fig. 5C, L) and *dlx5* (36.8%, Fig. 5F, L), whereas pax2 showed increased expression (30.0%, Fig. 5I, L). Comparing the frequencies of gene expression changes between embryos injected with wild type *sobp* versus *R651X* mRNAs showed effects on *six1* were similar (p=0.742), whereas *dlx5* and *pax2* were affected by R651X significantly less frequently (*dlx5*: p<0.0001; *pax2*: p<0.05). qPCR confirmed that the effects on otic gene expression were reduced in R651X-expressing embryos as there were no significant changes in *six1* (p=0.671981/q=0.678701), *dlx5* (p=0.621603/q=0.678701) or *pax2* (p=0.435624/q=0.678701) mRNA levels (Fig. 5O). Taken together, these data indicate that even though the R651X mutation mildly disrupts PPE patterning, it is less disruptive than full-length Sobp on otic vesicle gene expression.

### Sobp is required for craniofacial cartilage development

Because altered Sobp levels affected NC gene expression (Fig. 4) we examined whether this leads to defects in cranial cartilages at tadpole stages (Fig. 6). Gross analysis of surviving tadpole heads revealed *sobp* KD by CRISPR caused mild to severe cranial cartilage hypoplasia on the injected side in a subset (5nl: 18.6%; 10nl: 46.7%) of F0 tadpoles, whereas injection of *sobp* mRNA caused only mild hypoplasia (Fig. 6A-B). Tadpoles injected with *R651X* did not have apparent cranial cartilage defects (Fig. 6C). Staining tadpoles with Alcian Blue demonstrated that the observed hypoplasia in Sobp F0 crispants was the consequence of deformed Meckel’s and ceratohyal cartilages, hypoplastic branchial arch cartilages, and absent quadrate and otic cartilages (Fig. 6D, G, J). Most of the cartilaginous elements of the *sobp* mRNA-injected tadpole heads were normal; only the otic capsule was mildly hypoplastic in 56.6% of tadpoles (Fig. 6E, H, J). *R651X* mRNA-injected tadpole cranial cartilages, including the otic capsule, were minimally affected; only 3.8% of tadpoles showed a mildly hypoplastic otic capsule (Fig. 6F, I, J). These findings indicate that early changes in gene expression caused by loss or increased expression of Sobp lead to craniofacial skeletal defects, particularly disruption of the otic capsule.

**Figure 6.**
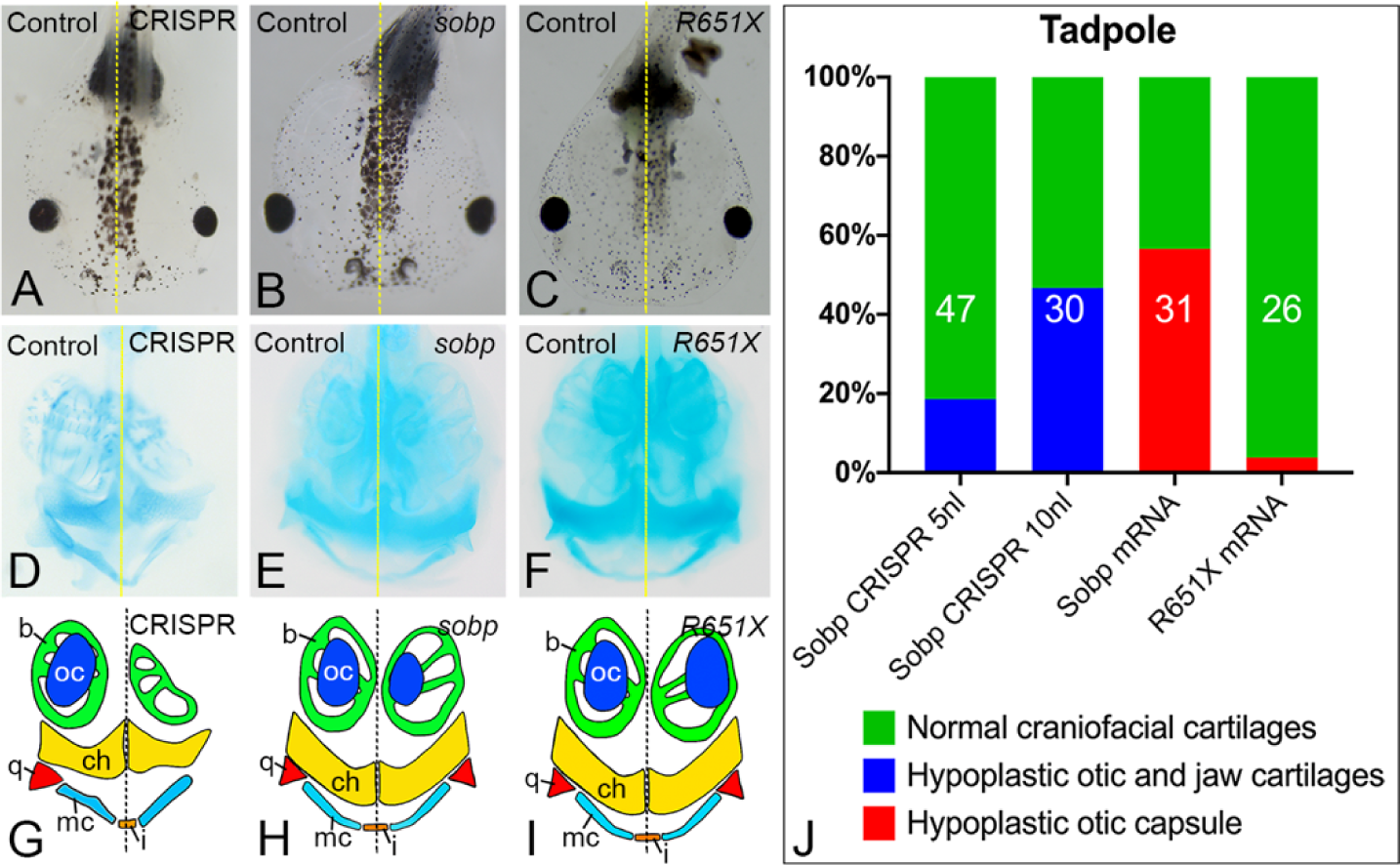
Sobp is required for craniofacial cartilage development. **A-C.** Gross morphology of tadpoles after unilateral (A) CRISPR knock-down (5nl), (B) increased *sobp* or (C) increased R651X. Survival rates were: CRISPR 5nl: ∼65.9%; 10nl: ∼40.0%; *sobp* mRNA: ∼96.3%; *R651X* mRNA: ∼87.2%. Hypoplasia of head structures on the injected side is noticeable from a dorsal view in a subset of Sobp Crispants and increased Sobp, but not of increased R651X. **D-I.** Ventral views of Alcian Blue staining of tadpoles (D-F) and drawings of the stained cartilages (G-I) show severe cranial cartilage defects of Crispants (5nl, 18.6%; 10nl, 46.7%), including deformed Meckel’s (mc) and ceratohyal (ch) cartilages, hypoplastic branchial arch cartilages (b), absent quadrate (q) and absent otic capsule (oc) cartilages. Increased *sobp* resulted in hypoplasia of the otic capsule (56.6%), whereas the majority of the *R651X* mRNA-injected tadpoles (96.2%) did not have apparent defects. i, infrarostral cartilage. **J.** Frequencies of defects in the cranial cartilages depicted in D-I. The number in each bar denotes the sample size.

## DISCUSSION

Development of the cranial sensory placodes is a complex process that requires different signaling pathways and expression of transcription factors that induce and/or repress gene expression (Saint-Jeannet and Moody, 2014, Moody and LaMantia, 2015). Previous work from our lab and others showed that Six1 functions at early stages of placode formation (Christophorou et al., 2009, Brugmann et al., 2004), during otic vesicle patterning (Zheng et al., 2003, Ozaki et al., 2004, Laclef et al., 2003), and hair cell formation in the inner ear (Zhang et al., 2017, Li et al., 2020, Ahmed et al., 2012).

One important characterisitc of Six1 factors that allows them to act in different contexts is their interaction with co-factors that modulate their transcriptional activities (Neilson et al., 2020, Neilson et al., 2017, Li et al., 2003, Li et al., 2020, Kenyon et al., 2005a, Kenyon et al., 2005b, Heanue et al., 1999, Guo et al., 2011, Brugmann et al., 2004, Ahmed et al., 2012). We identified Sobp as a potential Six1 co-factor based on findings in *Drosophila* (Neilson et al., 2010, Kenyon et al., 2005a), but little is known about its function in vertebrate development or the mechanisms by which it interacts with Six1.

Herein, we demonstrate that Sobp is a bone fide Six1 co-factor that can repress Six1+Eya1 transcriptional activation. In addition, we show that altering the levels of Sobp disrupts the patterning of embryonic ectodermal domains and otic vesicle genes (Fig. 7).

**Figure 7.**
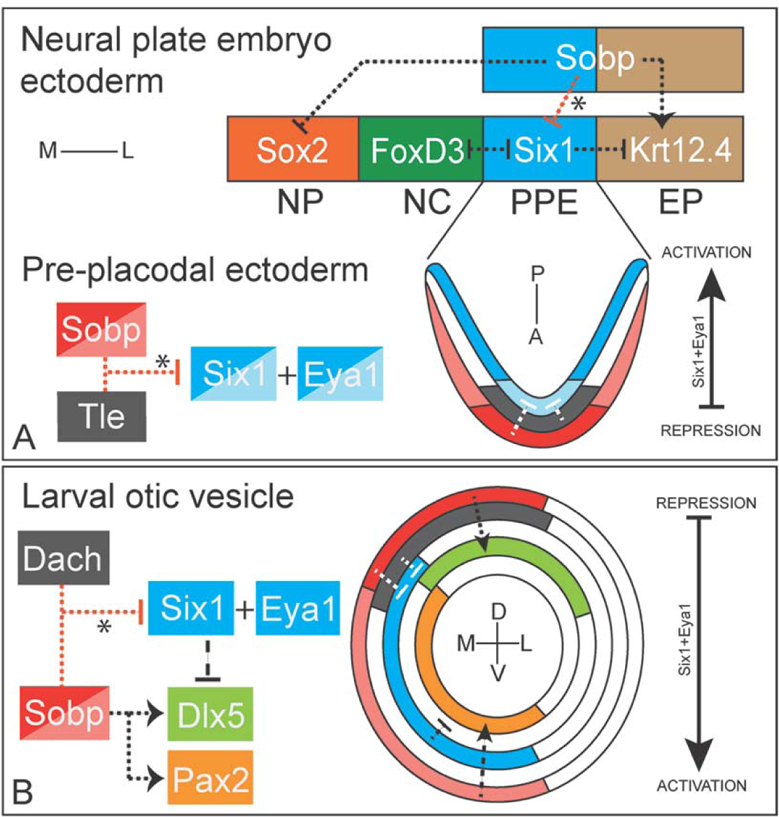
Model for Sobp interactions during craniofacial development. **A.** Sobp expression in two ectodermal domains (PPE and EP) directly or indirectly induces epidermal genes (*krt12.4*), represses neural plate genes (*sox2*) and represses the Six1+Eya1 transcriptional activation (*) in the PPE. Sobp effects outside the PPE are likely independent of Six1. Repression of Six1+Eya1 activation (*) is likely achieved by higher levels of Sobp and lower levels of Six1 and the additional expression of the known Six1 co-repressors (e.g., Groucho/Tle) in the anterior PPE. Conversely, in the posterior PPE, lower levels of Sobp, higher levels of Six1 and lower/no Groucho/Tle expression leads to transcriptional activation. Note that Eya1 is co-expressed with Six1 in this domain. A, anterior; EP, epidermis; L, lateral; M, medial; NP, neural plate; NC, neural crest; P, posterior; PPE, pre-placodal ectoderm. **B.** Sobp expression in the medial wall of the otic vesicle is higher in the dorsal region, a domain where Dach co-repressors are also expressed. This expression pattern leads to repression of Six1+Eya1 transcriptional activation (*), thus allowing expression of dorsal otic genes (e.g., *dlx5)*. Ventrally, Sobp induces expression of *pax2* independent of Six1. Six1 and Eya1 have overlapping expression. D, dorsal; L, lateral; M, medial; V, ventral.

### Sobp is a Six1 co-repressor

Sobp was first characterized in *Drosophila* as part of the retinal gene determination network based on its co-expression in the eye disc with So (Silver et al., 2003, Kenyon et al., 2005a). Our work is the first demonstration that in vertebrates Sobp interacts with Six1 and Eya1. Binding to Six1 was expected since *Drosophila* Sobp binds to So (Kenyon et al., 2005a), but binding to Eya1 was not expected; this interaction is reminiscent of fly Eya binding to the So co-repressor Dachshund (Chen et al., 1997). Similar to Eya factors, Sobp does not have DNA binding domains (Birk et al., 2010) and therefore relies on interaction with transcription factors to modulate transcription. However, unlike Eya1, a cytosolic protein that requires binding to Six factors for nuclear translocation (Ohto et al., 1999, Buller et al., 2001), Sobp is a nuclear protein (Chen et al., 2008, Birk et al., 2010). Our findings demonstrate that Sobp has a C-terminal NLS and after deletion of this critical domain or a larger C-terminal domain containing the NLS, Sobp is located almost exclusively in the cytosol. Interestingly, deletion of this NLS or the 220 C-terminal amino acids including the NLS (R651X mutant) still repressed Six1+Eya1 transcriptional activation after binding to Six1 and translocation to the nucleus. These data indicate that an interaction between Six1 and cytosolic proteins is not exclusive to Eya factors (Ohto et al., 1999).

Upon binding to the Six1+ Eya1 complex, Sobp reduces its ability to activate transcription. Our findings indicate that this repressive function occurs through a dose-dependent competition mechanism: high levels of Sobp disrupt the interaction between Six1 and Eya1 leading to at least some Eya1 remaining in the cytosol. The presence of FCS zinc finger domains (ZF1 and ZF2) (Chen et al., 2008) and SUMO interacting motifs (SIMs) (Sun and Hunter, 2012) of Sobp may be responsible for its repressive activity. The ZF domains are similar to those in constituents of the polycomb repressive complex 1 (PRC1) that modifies histones in chromatin leading to transcriptional repression (Chen et al., 2008, Blackledge and Klose, 2021). SUMOylation also causes transcriptional repression by modifying transcription factors and the assembly of nuclear protein complexes (Sun and Hunter, 2012), and SUMOylation of PRC1 is required for repression (Gill, 2010). Thus, the ZF and SIM domains in Sobp may account for its ability to alter the activity of the Six1+Eya1 complex and lead to repression. This needs to be tested experimentally via further mutations, but is consistent with our finding that R651X, which contains the ZF and SIM domains, repressed Six1+Eya1 transcriptional activation. It is interesting that BOR and some MRAMs patients present hearing loss but there are no reports of defects in balance caused by vestibular perturbations (Smith, 2018, Birk et al., 2010, Basel-Vanagaite et al., 2007), whereas the *jc* mouse presents with both hearing and balance defects (Chen et al., 2008, Calderon et al., 2006). A comparison between the MRAMs R661X and the *jc* S449fsX490 shows that the latter loses additional domains including both SIMs and a portion of the proline rich domain (PRD). Since PRDs can be involved in protein/protein interactions (Yu et al., 1994), the *jc* mutation could interrupt binding to Six1 or other factors. It will be important to test these various aspects in *jc sobp* to further understand protein structure-function relationships.

### Sobp is required to establish embryonic ectodermal domains

The PPE gives rise to all cranial sensory placodes and is characterized by the expression of *six1* and *eya1* (Schlosser and Ahrens, 2004, Pandur and Moody, 2000). *sobp* also is expressed in the PPE (Neilson et al., 2010), but we noticed by ISH that its expression is weaker in the posterior placodal region that will give rise to the otic and epibranchial placodes. This regional difference in expression is supported by a single cell RNA-seq (scRNA-seq) study that allowed us to compare the anterior and posterior placode regions of neural plate stage embryos (Briggs et al., 2018, Peshkin and Kirschner, 2020). *sobp* enrichment occurs in the anterior placodal cells that co-express *tle1* and *tle2*, two members of the Groucho family of known Six1 co-repressors (Roth et al., 2010). Together, these findings suggest that at PPE stages, Six1 may induce gene expression in the posterior PPE by transcriptional activation and its activity may be limited in the anterior PPE by the presence of multiple co-repressors including Sobp (Fig. 7A). This is consistent with the proposed role of Sobp in the fly eye field (Kenyon et al., 2005a).

The changes in gene expression after loss or increased Sobp expression that we observed also show that Sobp is required for the proper balance in size of the other embryonic ectodermal domains: neural plate, neural crest and epidermis. Our data indicate that Sobp appears to have a role in inducing directly or indirectly epidermal genes and repressing neural ectodermal genes (Fig. 7A). It is likely that NC and PPE were disrupted after loss of Sobp because the neural plate domain expanded, and that they were disrupted after increasing Sobp levels because the epidermis expanded. This function appears to be independent of its ability to reduce Six1 transcriptional activation because Six1 is not expressed in these other domains (Pandur and Moody, 2000, Briggs et al., 2018, Peshkin and Kirschner, 2020). scRNA-seq analyses show that at neural plate stages *sobp* is not expressed in cells expressing *sox2* but is co-expressed in *krt19+* cells (Briggs et al., 2018). Because increased expression of the truncated R651X caused similar changes compared to full-length Sobp in the PPE but not to genes in the other domains, we posit that the truncation in *Xenopus* R651X/human R661X does not disrupt the interaction with Six1. It is, however, possible that the truncation hampers interactions with other nuclear factors because R651X cannot translocate into the nucleus in domains where Six1 is not present. It will be important to determine with which other proteins Sobp interacts in different embryonic ectodermal domains.

### Sobp contributes to patterning the otic vesicle

Our *in vivo* knock-down studies show that Sobp does not appear to be required for *six1* expression in the otic vesicle. The decrease in otic *six1* after increasing either Sobp or R651X is likely due to a reduction in the interaction between Six1 and Eya1 since in mouse Eya1 is required for otic vesicle expression of *Six1* (Zheng et al., 2003). Although changes in gene expression in the otic vesicle may be influenced by earlier changes in the PPE, because *six1* is still expressed in the majority of Sobp crispants, changes in PPE gene expression do not completely account for otic vesicle changes.

Instead, we propose that the Sobp-Six1 relationship in the otic vesicle likely impacts dorsal-ventral (D-V) patterning.

D-V patterning of the otic vesicle is critical for the development of ventral auditory structures and dorsal vestibular structures (Ohta and Schoenwolf, 2018, Nakajima, 2015, Bever et al., 2003). Published data and data presented here show that Six1, Eya1 and Sobp are co-expressed in the medial wall of the otic vesicle (Durruthy-Durruthy et al., 2014, David et al., 2001). Unlike the apparent uniform expression of *six1*, *sobp* expression is more intense dorsally, a region that in mouse express two known Six1 co-repressors: *Dach1* and *Dach2* (Ozaki et al., 2004, Li et al., 2002, Li et al., 2003, Durruthy-Durruthy et al., 2014). These expression data suggest that Six1+Eya1 transcriptional activation is repressed in the dorsal-medial domain of the otic vesicle, in agreement with previous findings in mouse that Six1 and Eya1 induce expression of ventral genes such as *Otx*1 while repressing dorsal genes such as *Dlx5* (Zheng et al., 2003, Ozaki et al., 2004). We suggest that in the otic vesicle there is likely a dose-dependent mechanism of Six1 activation and repression involving co-factors such as Eya1, Sobp and Dach that contributes to D-V patterning (Fig. 7B). Dorso-medially, higher levels of Sobp and Dach would repress Six1, allowing localized expression of dorsal genes such as *dlx5.* Ventro-medially, Sobp induces *pax2* expression independent from Six1, since *Pax2* expression does not depend on Six1 in mouse (Ozaki et al., 2004); concurrently, Six1 and Eya1 are transcriptionally active in this domain because of lower levels of Sobp and absence of Dach. Sobp likely interacts with other factors to accomplish otic D-V patterning since, compared to full-length Sobp, changes in gene expression were less frequent after increased R651X, which required increased expression of Six1 for nuclear translocation.

### Sobp effects on cranial cartilages

While we have focused our analyses on early stages of otic development, the changes after altering Sobp levels in the embryonic ectodermal domains ultimately led to severe cranial cartilage defects. The differences in severity between loss and increased Sobp expression might be explained by the targeting approach: for CRISPR/Cas9-mediated KD, embryos were injected at the 2-cell stage, whereas mRNA injections were performed at the 8-cell stage only in precursors of the PPE and NC. Because Sobp appears to not be expressed in *Xenopus* branchial arches (Neilson et al., 2010), the defects in the cranial cartilages likely are secondary to changes in the early ectodermal domains rather than disruption of NC patterning in the branchial arches. Although additional work is required to understand Sobp function during formation of these cartilages, it is notable that the otic cartilage was consistently hypoplastic when levels of Sobp were altered. Perhaps this defect contributes to the hearing and vestibular deficits observed in *jc* mice.

In summary, we show that Sobp is a Six1 co-repressor that interacts with Six1 and most likely other factors during otic and craniofacial development. Although Sobp does not have an identified DNA binding domain (Birk et al., 2010), it functionally modulates the Six1-Eya1 transcriptional complex. The mutations found in MRAMs patients (Birk et al., 2010) and *jc* mice (Chen et al., 2008) that have overlapping but also very distinct phenotypes suggest that disruption of different domains in Sobp may perturb interactions with different partners. To better understand the function of Sobp during normal development and in congenital syndromes, it will be essential to assess the role of its various domains and identify any additional binding partners.

## MATERIALS AND METHODS

Many of the methods were supported by Xenbase (http://www.xenbase.org/, RRID: SCR_003280) and the National *Xenopus* Resource (http://mbl.edu/xenopus/, RRID:SCR_013731).

### Plasmid constructs

A full-length *Xenopus tropicalis sobp* plasmid was purchased from Open Biosystems (BC154687); subsequent sequence analysis identified it as having ∼95% homology to the predicted sequences for *Xenopus laevis* L-homeolog sobp (XM_018263336.1; XM_018263335.1) and S-homeolog sobp (XM_018265405.1). The ORF was subcloned into the BamH1 site of *pCS2^+^* (*pCS2^+^-sobp*) using the Clone EZ PCR cloning kit (GenScript). *pCS2^+^-5’HA-sobp* and *pCS2^+^-sobp-3’HA* were generated using the QuikChange lightning Site-directed mutagenesis kit (Agilent). The same kit was used to sequentially remove three nucleotides located at the 5’ end of the *pCS2^+^-3’HA-sobp* ORF to generate a construct whose transcribed mRNA does not bind to the designed translation-blocking antisense morpholino oligonucleotide (*pCS2^+^-sobpMOI-3’HA*; morpholino insensitive). To generate a plasmid containing the human R661X mutation, the mutagenesis kit was used to introduce into the ORF of *pCS2^+^-sobp* and of *pCS2^+^-5’HA-sobp* a stop codon at aa 651 (c.1951A<T; p.R651X; *pCS2^+^-R651X* and *pCS2^+^-5’HA-R651X)*. Finally, a conserved putative C-terminal nuclear localization signal (NLS) was identified by analyzing the amino acid sequences for *Xenopus tropicalis* Sobp (NP_001096678.1), *Xenopus laevis* Sopb (XP_018118825.1; XP_018120894.1), Homo sapiens SOBP (NP_060483.3) and *Mus musculus* Sobp (NP_780616.4) using cNLS mapper (http://nls-mapper.iab.keio.ac.jp/cgi-bin/NLS_Mapper_form.cgi). The putative NLS was deleted using the mutagenesis kit (*pCS2^+^-5’HA-sobp-NLSdel).* All plasmids were confirmed by full-length sequencing in both directions.

### Cell transfection

HEK293T/17 cells (ATCC CRL-11268) were grown in Dulbecco’s modified Eagle medium (Hyclone) supplemented with 10% fetal bovine serum (Lonza) and penicillin-streptomycin (Gibco). Cells were plated into 24-well plates (Fisher) for luciferase assays; into 6-well plates (Fisher) for co-immunoprecipitation experiments (Co-IP); and into 1-well Nunc Lab-Tek Chamber slides (Thermo) for immunofluorescence (IF). Cells were transfected according to the manufacturer’s protocol using X-tremeGENE 9 DNA (Sigma) for luciferase assays and IF; and lipofectamine 3000 (Thermo) for Co-IP. Cells were processed for each assay 48h after transfection.

### Co-Immunoprecipitation

HEK293T cells were transfected with combinations of *pCS2^+^-3’Flag-Six1*, *pCS2^+^-5’HA-Sobp, pCS2^+^-5’HA-sobp-NLSdel, pCS2^+^-5’HA-R651X* and/or *pCS2^+^-5’Myc-Eya1*. 1µg of each plasmid was used unless noted in the figure legend. Cells were extracted after 48h using 500µl of ice-cold Pierce IP lysis buffer (Thermo) plus Halt Protease Inhibitor Cocktail with EDTA (Thermo). Cell debris was pelleted at 13,000g for 10 min at 4°C.

The supernatant was subjected to immunoprecipitation using the Pierce anti-HA magnetic beads (Thermo) or the Pierce anti-DYKDDDDK magnetic agarose (Thermo). Proteins were washed five times and eluted using the Pierce lane marker non-reducing sample buffer (Thermo). The eluted proteins were reduced using 100mM DTT at 100°C for 5 min prior to SDS-PAGE and Western blot. For control experiments, 10µl of each sample in IP lysis buffer was diluted with Laemmli sample buffer with 2% BME, incubated at 100°C for 5 min prior to SDS-PAGE and Western blot. Immobilon-FL PVDF membranes (Fisher) were probed with mouse anti-HA (6E2, Cell Signaling, 1:1,000) to detect Sobp, mouse anti-Myc (9B11, Cell Signaling, 1:1,000) to detect Eya1, rabbit anti-Six1 (D5S2S, Cell Signaling, 1:1,000) and rabbit anti-β-actin (13E5, Cell Signaling, 1:1,000). Secondary antibodies were IRDye 680RD donkey anti-rabbit IgG (92568073, Licor, 1:5,000) and IRDye 800CW goat anti-mouse IgG (92532210, Licor, 1:5,000).

Experiments were repeated at least three independent times. Blots were scanned using the Licor Odyssey infrared imaging system.

### Luciferase assay

Transfected HEK293T cells were harvested and analyzed using the Dual Luciferase Assay kit (Promega) following the manufacturer’s directions. Each transfection included 200 ng of *pGL3-6XMEF3-Firefly luciferase* reporter (Ford et al., 2000) and 100 ng of *Renilla* luciferase reporter (*pRL-TK*), in addition to different combinations of pCS2^+^ (control), pCS2+-3’Flag-Six1, pCS2+-5’Myc-Eya1, pCS2+-5’HA-Sobp, pCS2+-5’HA-sobp-NLSdel and/or pCS2+-5’HA-R651X (400ng each). At 48h post-transfection, cells were resuspended directly in 100 µl of passive lysis buffer (PLB) and 20 µl of lysate was used in the analysis. Experiments were repeated at least five times. ANOVA with Tukey post hoc multiple comparisons test was performed using GraphPad Prism 9 software.

Expression of exogenous proteins from the transfected plasmids was confirmed by standard Western blotting using antibodies described for Co-IP (Fig. S2).

### Immunofluorescence

HEK293T cells were transfected with different combinations of *pCS2^+^-3’Flag-Six1*, *pCS2^+^-5’HA-Sobp*, *pCS2^+^-5’HA-sobp-NLSdel*, *pCS2^+^-5’HA-R651X* and/or *pCS2^+^-5’Myc-Eya1*. 2µg of each plasmid was used unless noted in the figure legend. Cells were processed as described previously (Shah et al., 2020). Briefly, 48 hours after transfection, cells were fixed in 4% paraformaldehyde and processed for immunostaining by standard methods using mouse anti-Flag (9A3, Cell Signaling, 1:400), rabbit anti-Myc (71D10, Cell Signaling, 1:250), rabbit anti-HA (C29F4, Cell Signaling) or mouse anti-HA (6E2, Cell Signaling, 1:800) followed by Alexa Fluor 488-conjugated anti-rabbit (4412, 1:1,000) and Alexa Fluor 568-conjugated anti-mouse (A1104, 1:1,000) secondary antibodies, and DAPI nuclear counterstain (R37605, Thermo). Experiments were repeated at least three independent times, and at least five fields per slide analyzed using a Zeiss LSM 710 confocal microscope.

### In vitro synthesis of mRNAs and antisense RNA probes

mRNAs encoding *Xenopus tropicalis sobp*, *Xenopus tropicalis 5’HA-sobp*, *Xenopus tropicalis sobpMOI-3’HA*, *Xenopus tropicalis R651X* and a nuclear-localized β *galactosidase* (*n gal*) lineage tracer were synthesized *in vitro* according to β manufacturer’s protocols (mMessage mMachine kit, Ambion). Antisense RNA probes for in situ hybridization (ISH) were synthesized *in vitro* (MEGAscript kit; Ambion), as previously described (Yan et al., 2009b).

### Morpholino oligonucleotide knock-down

To knock-down endogenous levels of Sobp protein in embryos, a 3’-carboxyfluorescein-labelled translation-blocking antisense morpholino oligonucleotide (MO) was purchased (GeneTools, LLC): TCCCCTCTTTTTCCATTTCTGCCAT. The MO binds at the ATG start site for both the predicted L-and S-homeologs for *Xenopus laevis* (Fig. S4A) and to *Xenopus tropicalis sobp*. To verify the ability of the MO to block *sobp* translation (Fig. S4B), *Xenopus* stage VI oocytes were injected with 9ng of MO, and then injected with either 2ng of *5’HA-Sobp* (MO-insensitive), *Sobp-3’HA* (MO-sensitive) or *sobpMOI-3’HA* (MO-insensitive) mRNA. Oocytes were cultured overnight at 18°C. Lysates were prepared and Western blotting performed with rabbit anti-HA antibody (C29F4, Cell Signaling) as previously described (Neilson et al., 2017).The specificity of the MO was tested by injecting embryos with 9ng of MO followed by the immediate microinjection of 50pg of *sobpMOI-3’HA* mRNA, and processing embryos for *foxd3* expression by ISH (Fig. S4C-D), as described below.

### CRISPR/Cas9 gene editing

Based on the *Xenopus laevis sobp* genomic sequence obtained from Xenbase (http:/xenbase.org), a 20bp target was designed by CRISPRscan (http://www.crisprscan.org) during the *Xenopus* Genome Editing Workshop at the National *Xenopus* Resource at the Marine Biological Laboratory (Woods Hole, MA, USA). Potential off-targets were predicted by GGGgenome (https://gggenome.dbcls.jp) using the most up-to-date version of the *Xenopus laevis* genome. The 5’ dinucleotides were converted to GG (Gagnon et al., 2014). The sequence is GGTTCTTGGATGGTACGGTA and targets the L-and S-homeologs for *sobp* in its second exon (Fig. S4A) that encodes the first conserved region (Fig. S4A) (Chen et al., 2008, Kenyon et al., 2005a). A DNA template was produced by a PCR-based method using a universal reverse primer with a gene-specific forward primer containing a T7 polymerase promoter. The MEGAshortscript T7 transcription kit (Thermo) was used for sgRNA synthesis. sgRNA was mixed with Cas9 Protein (PNA Bio) and Texas Red Dextran, Lysine Fixable (Thermo) prior to injections. Injected embryos were incubated at 21°C and at least 8 embryos were processed for sequencing to confirm DNA editing for the L-and S-homeologs for *sobp* each time injections were performed. Genotyping primer sequences are listed in Table S1. Insertion/deletion frequencies were calculated with the TIDE software package (http://shinyapps.datacurators.nl/tide/; Fig. S4E-F).

### Embryo microinjections

Fertilized *Xenopus laevis* embryos were obtained by natural mating and *in vitro* fertilization (Moody, 2000). Injections for ISH and cartilage staining were performed unilaterally with 1 nl of MO (9 ng/nl) or 1nl of mRNA (100 pg mixed with 100 pg *n gal* β mRNA) in the dorsal-animal and ventral-animal blastomeres of 8-cell stage embryos; these blastomeres predominantly give rise to the neural crest and cranial placodes (Moody and Kline, 1990); or with 5 nl or 10 nl of 75pg/nl of sgRNA mixed with 0.2 ng/nl of Cas9 protein (PNA Bio) and 0.1% Texas Red Dextran in 1 cell of 2-cell stage embryos. Injections for quantitative real time PCR (qPCR) were performed with 1nl of MO (2 ng/nl) or 2nl of mRNA (100pg mixed with 100 pg *nβgal*) in each cell of 2-cell β stage embryos; or with 10 nl of sgRNA/Cas9 solution in 1-cell stage embryos. Microinjections were performed according to standard methods (Moody, 2018). Embryos were cultured in diluted Steinberg’s solution until fixation or harvest.

### Histochemistry and in situ hybridization

Embryos were cultured to neural plate (st. 16-18) and larval (st. 28-34) stages (Nieuwkoop and Faber, 1994), fixed in 4% paraformaldehyde (PFA), stained for β-Gal histochemistry, and processed for *in situ* hybridization (ISH) as described previously (Yan et al., 2009a). In embryos in which the fluorescent label/dextran or *n*β*gal* lineage tracer were located in the appropriate tissue domains, the position, intensity and size of the expression domains of *sox2*, *foxd3, six1, krt12.4, dlx5* and *pax2* were compared between the injected, lineage-labeled side to the control, uninjected side of the same embryo, thus accounting for inter-embryo variation. Embryos for each assay were derived from a minimum of three different sets of outbred, wild type parents. Differences in the frequency of gene expression changes were assessed for significance (p<0.05) by the Chi-square test using GraphPad Prism 9. A set of control larvae that was processed for *six1* and *sobp* ISH was embedded in a gelatin-based medium (0.5% gelatin, 30% bovine serum albumin, 20% sucrose, hardened with glutaraldehyde [75μl/ml]), and vibratome sectioned at 50 µm in the transverse plane. Whole-mount and serial section ISH images were collected with an Olympus SZX16 stereomicroscope coupled to an Olympus UC90 camera and cellSens Entry software.

### RNA collection and qPCR

Three embryos at neural plate (st. 16-18) or three dissected heads at larval (st. 28-34) stages were collected in TRI-reagent (Zymo) and processed for RNA extraction with DNAse I treatment using the Direct-zol RNA Miniprep kit (Zymo). cDNA was synthesized using the iScript Advanced cDNA Synthesis kit (Bio-Rad). qPCR was performed using 5ng cDNA with the SsoAdvanced Universal SYBR Green Mix (Bio-Rad). Primer sequences are listed in Table S2. qPCR of four biological replicates was performed in duplicate. PCR and data analysis were performed using a CFX Connect thermocycler (Bio-Rad). Statistical analysis was performed with GraphPad Prism 9, with significance calculated by unpaired t-tests followed by a False Discovery Rate approach using the two-stage step-up method (Benjamini et al., 2006)(FDR=1%).

### Cartilage staining

Embryos were grown to tadpole stages and the ones that remained alive were photographed and counted for quantification of survival rates. They were subsequently processed as described previously (Young et al., 2017): fixed in 4% PFA for 1 hour at room temperature then incubated in a solution of acid/alcohol containing 0.1% Alcian Blue. When staining was complete, tadpoles were washed in the acid/alcohol solution without Alcian Blue, bleached with a solution containing 1.2% hydrogen peroxide and 5% formamide and cleared in 2% KOH with increasing concentrations of glycerol.

## ACKNOWLEDGMENTS

This work was made possible with the support of Xenbase (http://www.xenbase.org/, RRID: SCR_003280) and the National *Xenopus* Resource (http://mbl.edu/xenopus/, RRID:SCR_013731). We thank the *Xenopus* Genome Editing Workshop at the National *Xenopus* Resource that helped with sgRNA design and training on CRISPR/Cas technology. We also thank Francesca Pignoni, Kristy Kenyon and Dominique Alfandari for helpful discussions during the execution of this work. Finally, we thank many members of the *Xenopus* community for providing plasmids. This work was supported by the National Institutes of Health (DE022065 to S.A.M., DE026434 to S.A.M and K.M.N).

---

**Figure S1.**
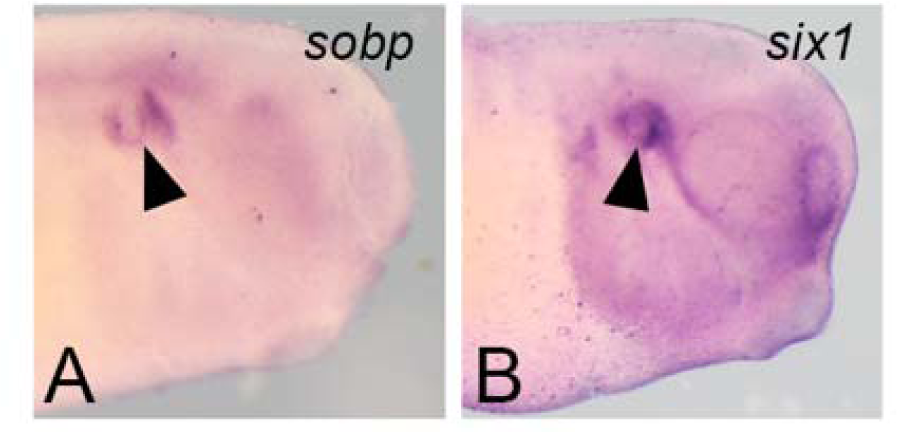
In situ hybridization for *sobp* and *six1* at larval stages. Whole-mount view of embryos sectioned in Fig. 1C (*sobp*) and D (*six1*) showing expression in the otic vesicle (arrowheads)

**Figure S2.**
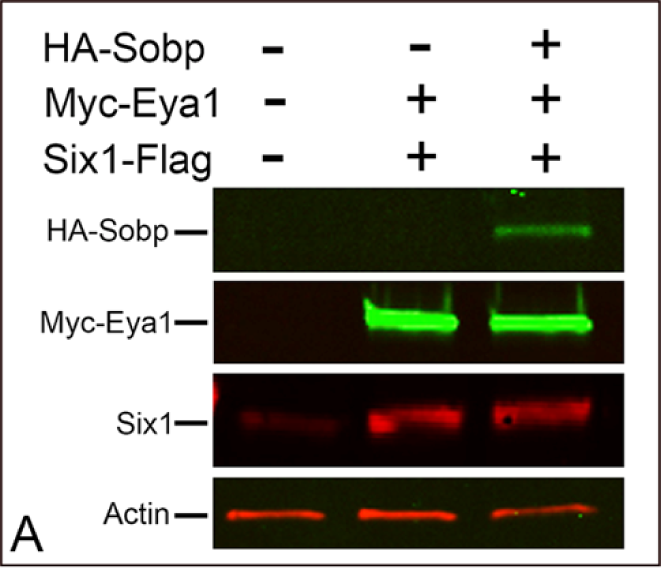
Control multiplex fluorescence Western blot for luciferase assays. Constructs for HA-Sobp, Myc-Eya1 and Six1-Flag are properly expressed in HEK293T cells in different combinations tested in luciferase assays. Actin is used as loading control.

**Fig. S3.**
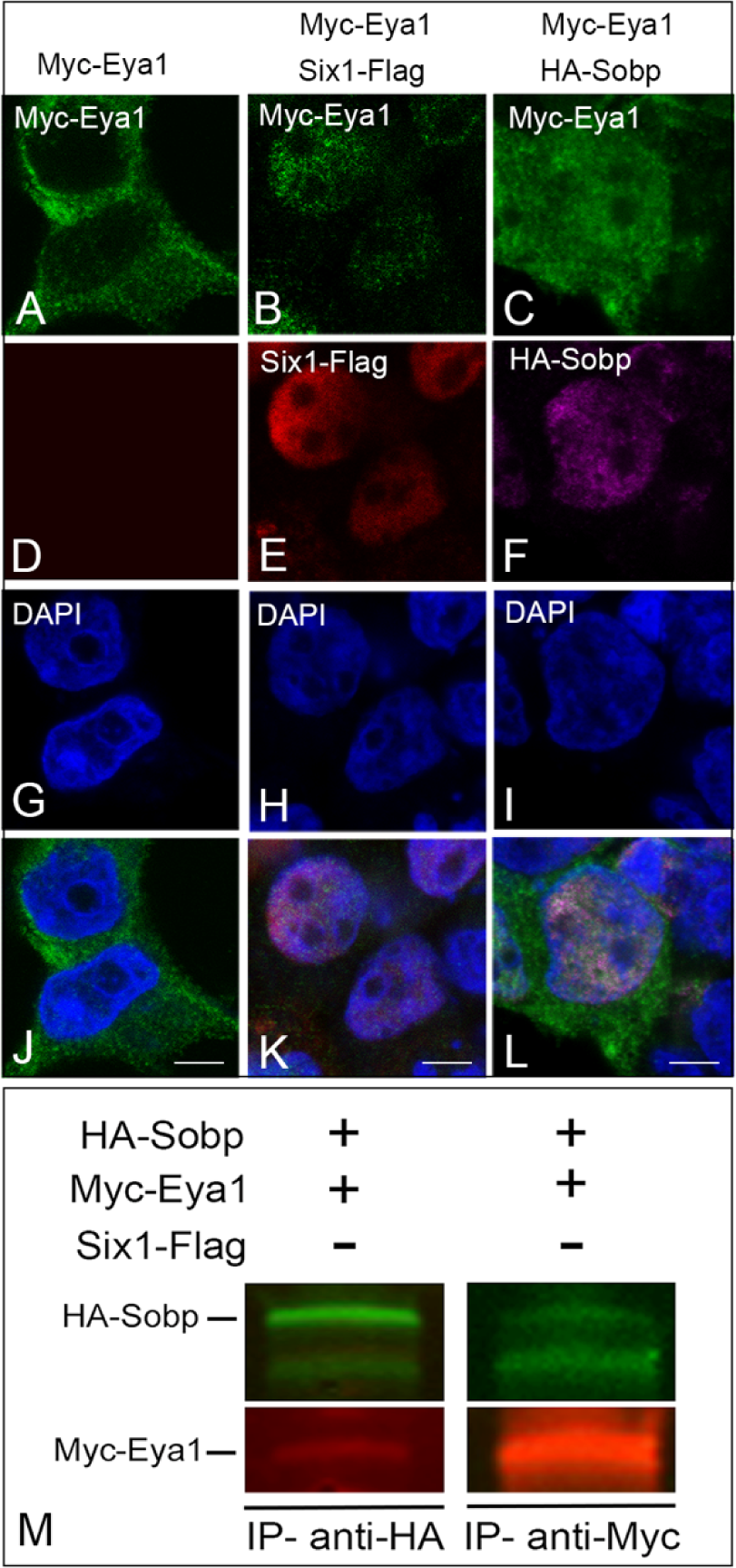
Sobp and Eya1 interact in the absence of Six1. **A-L.** Confocal images of HEK293T cells expressing Myc-Eya1 (green), Six1-Flag (red) and/or HA-Sobp (magenta). Myc-Eya1 is located exclusively in the cytosol (A, J) and is completely translocated to the cell nucleus by Six1-Flag in the majority of the cells (B, E, K). Surprisingly, HA-Sobp also partially translocates Myc-Eya1 to the cell nucleus (C, F, L) in the absence of Six1. Cell nuclei are stained with DAPI (blue in G-I). Bars: 5μm. **M.** HEK293T cells were co-transfected with HA-Sobp and Myc-Eya1 followed by multiplex fluorescence Western blot detection for HA-Sobp (green) and Myc-Eya1 (red). Myc-Eya1 is detected when HA-Sobp is immunoprecipitated (IP, anti-HA, left column). The reverse IP (anti-Myc, right column) confirmed this interaction.

**Figure S4.**
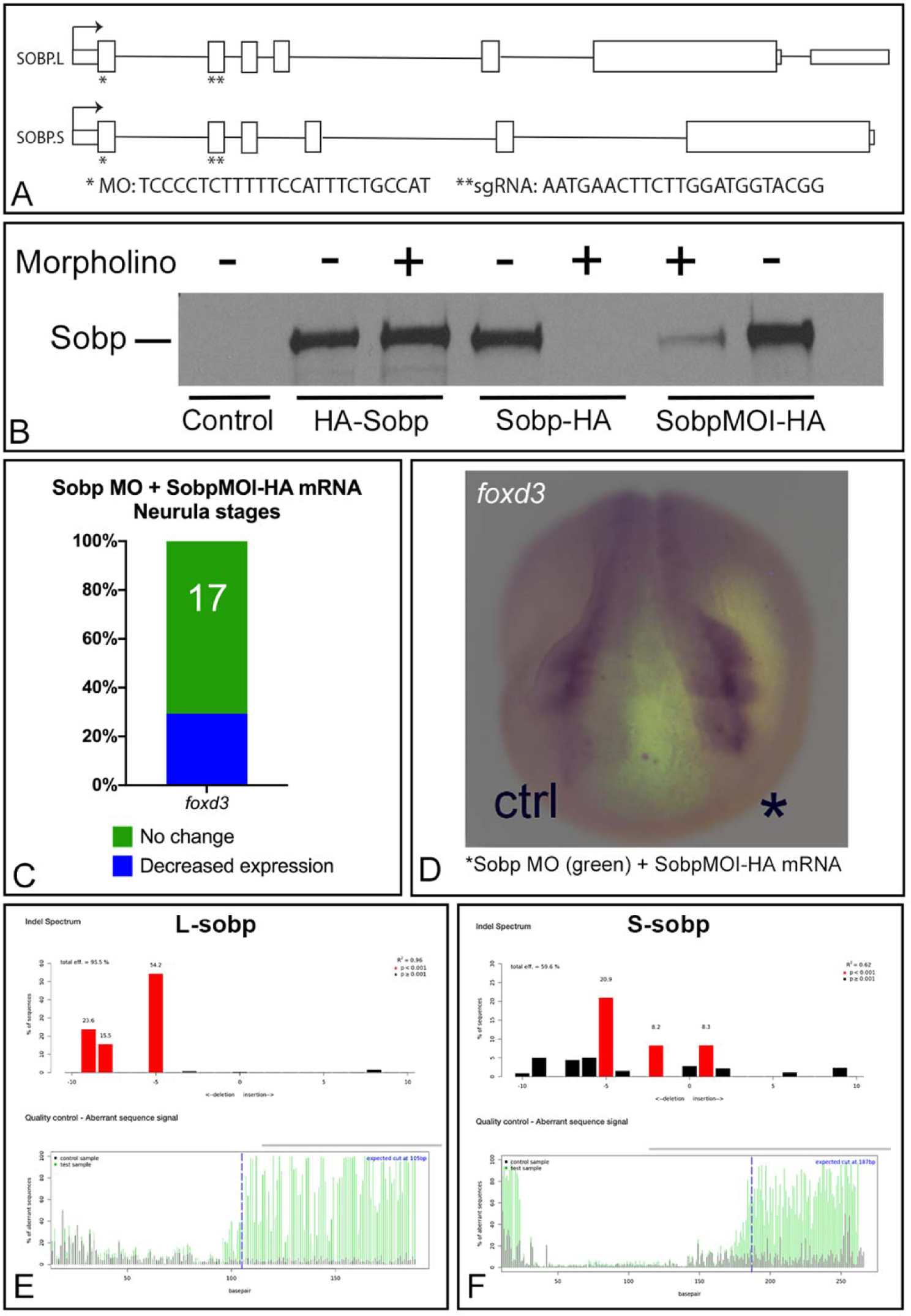
Control experiments for *in vivo* studies using a translation-blocking antisense morpholino oligonucleotide (MO) against Sobp or F0 analysis after CRISPR/Cas9-mediated genome editing. A. Schematic representation of exons and introns in *Xenopus laevis* Sobp.L and Sobp.S genes. The MO binds at the ATG start site for both L- and S-homeologs. The sgRNA targets the L- and S-homeologs in the second exon. **B.** Western blot detection of the HA tag showing the ability of the MO to block endogenous *sobp* translation (represented by Sobp-HA). The HA-*sobp* transcript is expected to avoid translation blockage because the 5’HA tag prevents MO binding at the translational start site. Translation of *sobp*-HA is expected to be blocked in the presence of the MO because the 3’HA tag does not interfere with MO binding. *sobpMOI*-HA is translated in the presence of MO because it has a deletion of the third codon in the *sobp* ORF (and a 3’HA tag) making it insensitive to a translation-blocking MO. **C.** Graph showing decrease in the frequency of *foxd3* reduction (29.4%) after partial rescue with unilateral injection of MO plus *sobpMOI*-HA mRNA (compare to 90.2% reduction in MO-only embryos). **D.** An embryo in which *foxd3* expression on the MO+*sobpMOI*-HA side (*) is similar to control side. **E-F.** Calculation of insertion/deletion frequencies with the TIDE software package in an injected embryo after CRISPR/Cas9 editing showing that sgRNA targets both homeologs at the predicted site.

**Table S1.**
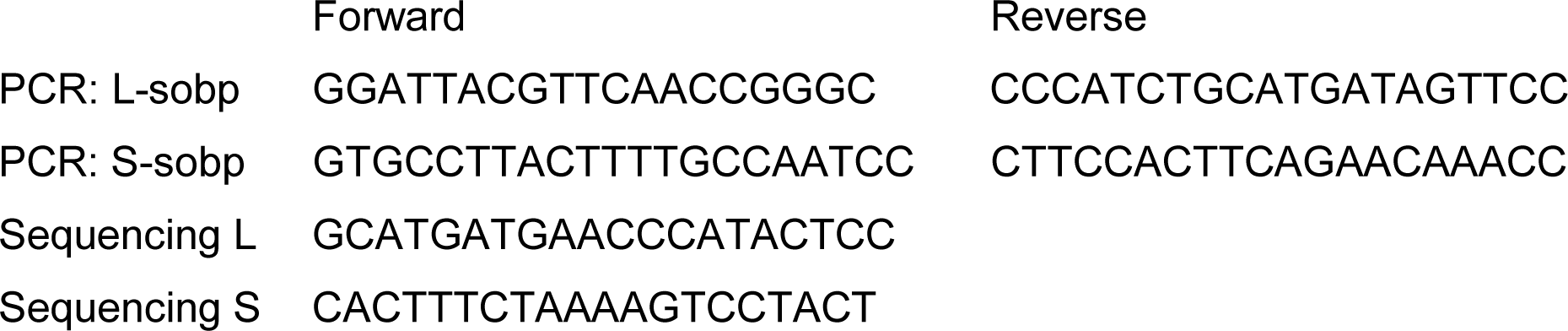
CRISPR primer and genotyping sequences

**Table S2.**
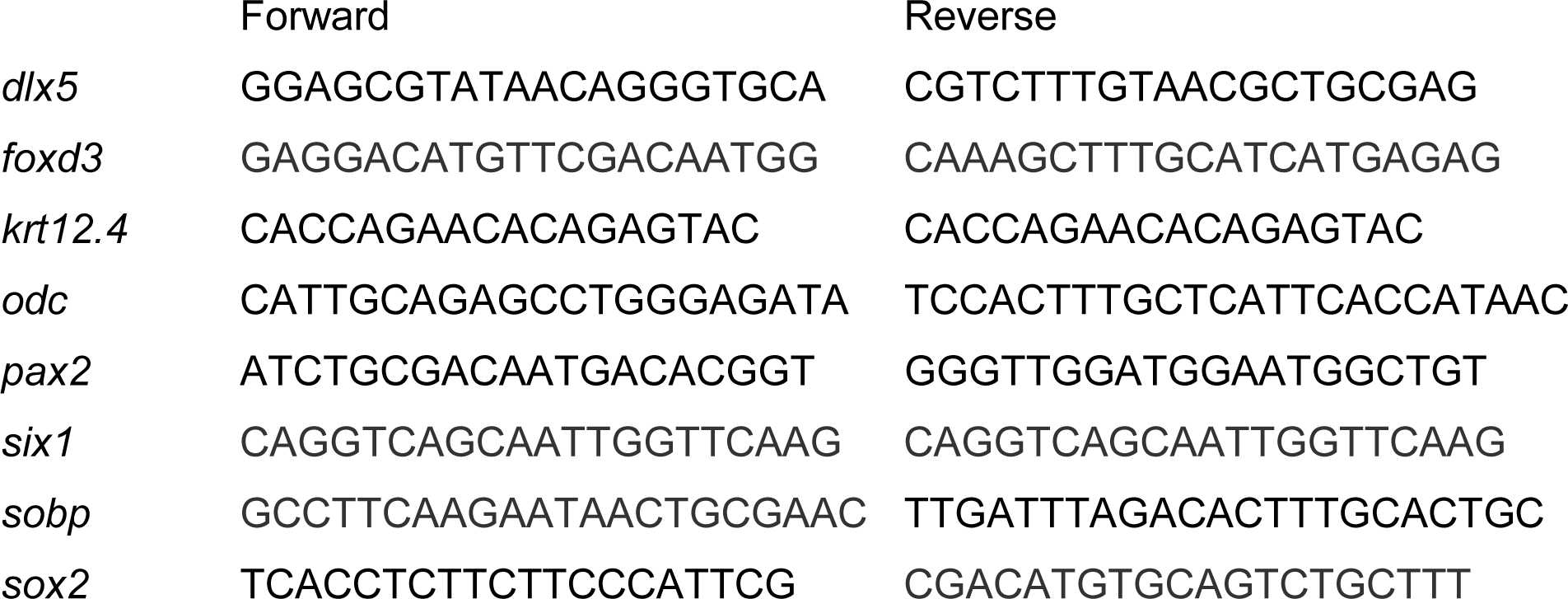
qPCR primer sequences

